# Tangential flow microfluidics for the capture and release of nanoparticles and extracellular vesicles on conventional and ultrathin membranes

**DOI:** 10.1101/675488

**Authors:** Mehdi Dehghani, Kilean Lucas, Jonathan Flax, James McGrath, Thomas Gaborski

## Abstract

Membranes have been used extensively for the purification and separation of biological species. A persistent challenge is the purification of species from concentrated feed solutions such as extracellular vesicles (EVs) from biological fluids. We investigated a new method to isolate micro- and nano-scale species termed tangential flow for analyte capture (TFAC), which is an extension of traditional tangential flow filtration (TFF). Initially, EV purification from plasma on ultrathin nanomembranes was compared between both normal flow filtration (NFF) and TFF. NFF resulted in rapid formation of a protein cake which completely obscured any captured EVs and also prevented further transport across the membrane. On the other hand, TFF showed capture of CD63 positive EVs with minimal contamination. We explored the use of TFF to capture target species over membrane pores, wash and then release in a physical process that does not rely upon affinity or chemical interactions. This process of TFAC was studied with model particles on both ultrathin nanomembranes and conventional thickness membranes (polycarbonate track-etch). Successful capture and release of model particles was observed using both membranes. Ultrathin nanomembranes showed higher efficiency of capture and release with significantly lower pressures indicating that ultrathin nanomembranes are well-suited for TFAC of delicate nanoscale particles such as EVs.

## Introduction

Tangential flow filtration (TFF) is used extensively in bioprocessing (van Reis & Zydney, 2001). In this method, a feed solution containing a species of interest flows tangentially over a selective membrane with some fraction of the flow also passing through the membrane. If the species of interest is to be retained behind the membrane, TFF can be used to remove impurities or to concentrate the species in the feed solution (Christy, Adams, Kuriyel, Bolton, & Seilly, 2002; Segura, Kamen, & Garnier, 2011). If the volume lost through transmembrane flow is resupplied to the feed channel as fresh buffer (diafiltration), TFF can be used for buffer exchange (Kurnik et al., 1995; Sweeny, Woehrle, & Hutchinson, 2006). TFF can also be used to partially purify a species that emerges in the filtrate (Fritsch & Moraru, 2008; van Reis et al., 1997), although the product typically requires final purification by column or membrane chromatography (Langfield, Walker, Gregory, & Federspiel, 2011; Lock, Alvira, & Wilson, 2012; Okada et al., 2009).

The advantage of TFF over normal flow filtration (NFF) is that the tangential flow component disrupts the formation of a concentration polarization layer that builds as species are rejected by the membrane (Belfort, Davis, & Zydney, 1994). Without a tangential component, this polarization layer will eventually form a ‘cake’ layer on the membrane with its own separation properties and significantly reduced permeate flux (Ghosh, 2006). With TFF filtration however, it is possible to identify conditions for which both the flux and transmembrane pressure (TMP) are steady with time (Field, Wu, Howell, & Gupta, 1995). Under these conditions filtration can, in principle, continue indefinitely.

Our laboratories develop ultrathin porous membranes for a range of applications including separations (Gaborski et al., 2010; Johnson et al., 2013; K. J. P. Smith, May, Baltus, & McGrath, 2017; Snyder et al., 2011). Ultrathin membranes are best defined as materials with pores on the same order as, or larger than, the membrane thickness (Mireles et al., 2017). These have been made with a variety of materials including silicon (Striemer, Gaborski, McGrath, & Fauchet, 2007), silicon-nitride (J. P. S. DesOrmeaux et al., 2014; Harris & Shuler, 2003; Vlassiouk, Apel, Dmitriev, Healy, & Siwy, 2009), silicon dioxide (Mazzocchi, Man, DesOrmeaux, & Gaborski, 2014), graphene (Surwade et al., 2015), and graphene-oxide (Nair, Wu, Jayaram, Grigorieva, & Geim, 2012). We have recently demonstrated that the high permeability of ultrathin membranes causes them to foul rapidly in NFF, with initial pore blockage events quickly followed by cake filtration (Winans, Smith, Gaborski, Roussie, & McGrath, 2016). We showed the same fouling phenomena occurs with both particle (Winans et al., 2016) and protein (K. J. Smith, Winans, & McGrath, 2016) solutes when used in NFF.

To extend the capacity of ultrathin membranes in separations, we have recently examined their performance in TFF. Working with undiluted serum and nanoporous silicon nitride (NPN) membranes (J. P. S. DesOrmeaux et al., 2014), we made the surprising discovery, reported here for the first time, that 60 - 100 nm extracellular vesicles (EVs), are captured in the pores of ultrathin membranes with little evidence of protein fouling. Our discovery inspired a closer look at the mechanisms and potential utility of capturing nanoparticle-sized analytes from biofluids in the pores of ultrathin membranes.

Extracellular vesicles (EVs) are secreted from tissue cells into all body fluids, and EVs that are < 100 nm are typically, but not exclusively, exosomes. Exosomes contain the largest pool of extracellular RNA (exRNA) in biofluids (Fernando, Jiang, Krzyzanowski, & Ryan, 2017; Gallo, Tandon, Alevizos, & Illei, 2012), and are thus valued both for their diagnostic (Armstrong & Wildman, 2018; Lane, Korbie, Hill, & Trau, 2018; M. Li et al., 2014) and therapeutic potential (Conlan, Pisano, Oliveira, Ferrari, & Pinto, 2017). The conventional method for exosome purification is ultracentrifugation although many alternative strategies have been proposed (P. Li, Kaslan, Lee, Yao, & Gao, 2017), including TFF (Busatto et al., 2018; Kang, Oh, Ahn, Lee, & Moon, 2008; McNamara et al., 2018). Out of respect for the careful criterion used to define exosomes (Théry et al., 2018), we will refer to < 100 nm EVs as small EVs (sEVs) rather than exosomes.

We propose a novel method for the extraction of nanoparticle species from biofluids which we call tangential flow for analyte capture (TFAC). In this method, sEVs and similarly-sized analytes are captured in the pores of an ultrathin membrane where they can be washed and released with additional flows. TFAC resembles bind/elute purification strategies although it distinguishes itself from affinity chromatography because the binding is purely physical. TFAC does not require engineered surface chemistries for capture or chemical treatments for elution. The purpose of the current report is to demonstrate the basic principles of TFAC using model particles. We also test the hypothesis that ultrathin membranes are ideally suited for TFAC because they facilitate capture and release at lower pressures than conventional thick membranes.

## Materials and Methods

### Fabrication of NPN Membranes

The fabrication steps for nanoporous silicon nitride nanomembranes (NPN) have been published previously (J. P. DesOrmeaux et al., 2014). Briefly, a silicon wafer is coated with a three layer stack of silicon nitride (SiN), amorphous silicon, and silicon dioxide. A porous nanocrystalline silicon (pnc-Si) layer is formed on top of SiN via rapid thermal annealing. The nanopores present in the pnc-Si are transferred into the SiN layer by reactive ion etching. In order to create the freestanding membranes, the back side of the silicon wafer is etched to the silicon nitride layer using ethylene diamine pyrocatechol.

### NPN Device Fabrication

Polydimethylsiloxane (PDMS) sheets (Trelleborg Sealing Solutions Americas, Fort Wayne, IN) were used to create microfluidic devices. Custom ordered 100 μm and 300 μm thick restricted grade sheets were patterned using a Silhouette Cameo digital craft cutter (Silhouette America, Oren, UT) (Yuen & Goral, 2010). The patterned silicone sheets were assembled into layer stack devices by aligning the patterned layers (Figure 1A, **Supplementary Movie 1**). NPN membrane chips (300 μm thick) were sandwiched between stacked layers and the final device was clamped to seal it for flow.

**Figure 1:**
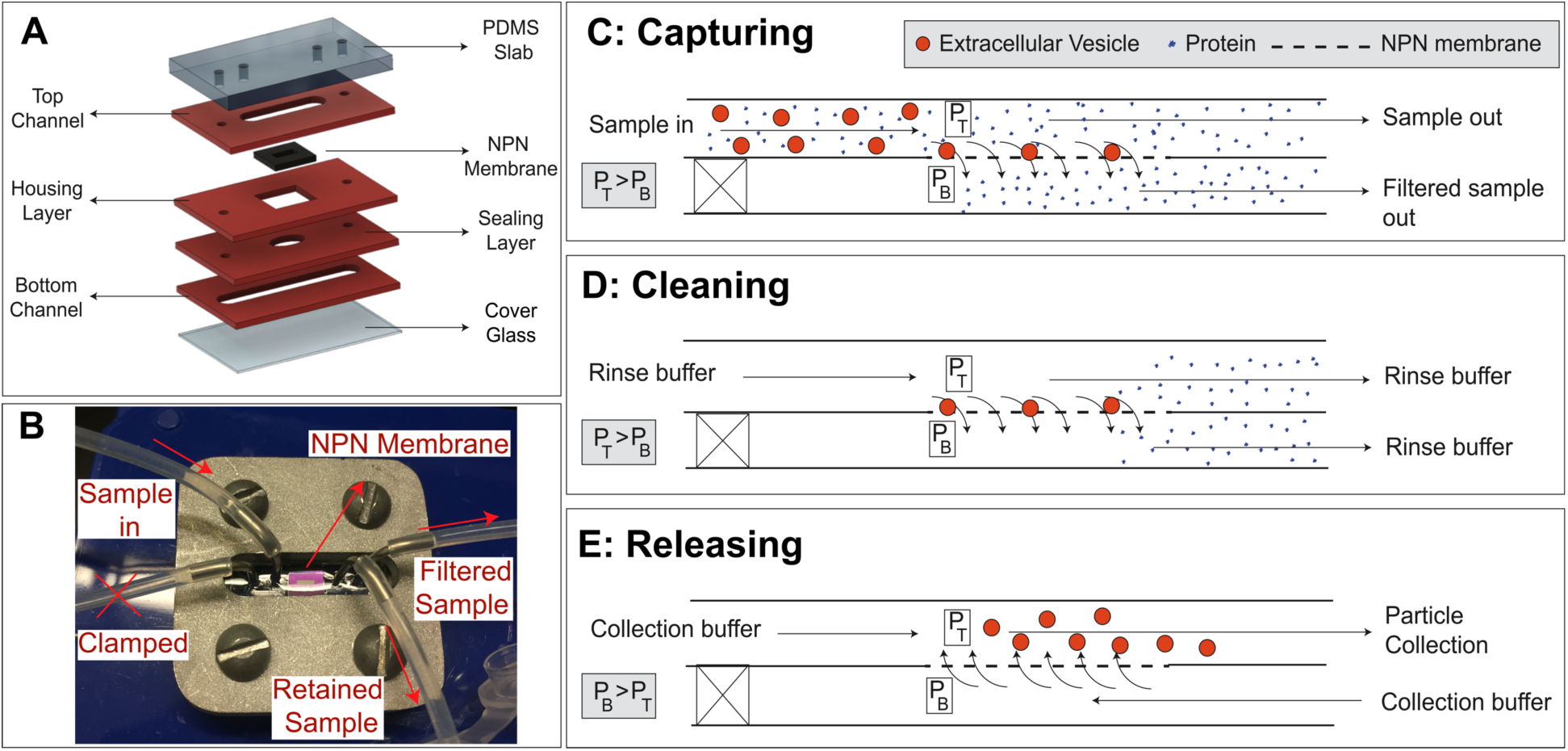
Tangential flow analyte capture (TFAC) technique for isolation of particles. **A)** Microfluidic devices are assembled through a layer stack process, in which channels and other featured are patterned into PDMS sheets. **B)** These layers are then formed into the device through thermal bonding or stacking and clamping. **C)** The sample is passed across the surface of the membrane and a transmembrane pressure generated by syringe pumps drives particle motion towards the membrane. Contaminating particles pass through pores or are swept downstream while the particles are retained on the membrane surface. **D)** The cleaning buffer is then passed through the input channel under the same flow condition as the capturing step to wash the channel and membrane surfaces of any remaining contaminants. **E)** The transmembrane pressure is then reversed, releasing the particles from the membrane where they are then swept downstream and collected.

### PCTE Device Fabrication

As a representative of conventional thickness membranes, commercial polycarbonate track-etch (PCTE) membranes with pore sizes of 8 μm and 80 nm were utilized (Sterlitech, WA, USA). In order to have a sealed system for track-etch membranes, the above described microfluidic device was modified. Holes were drilled in polycarbonate slabs for accessing the bottom channel of the device, while the PDMS slabs were punched for flowing to the top channel. We used 100 and 300 micron thick patterned PDMS sheets for bottom and top channels, respectively. In order to prevent leaking in the system, the PCTE membranes covering the entire device were sandwiched between the top and bottom layers using a clamp (Supplementary figure 1).

### sEV CAPTURE FROM PLASMA

#### Normal Flow Filtration

Small extracellular vesicle experiments were performed using purified human plasma (Equitech-Bio, Inc., Kerrville, TX). NFF experiments were performed using NPN chips with 50 nm thick freestanding membranes, with an average pore diameter of 50 nm and a porosity of 15% in a SepCon™ centrifuge cup (SiMPore Inc., Rochester, NY). A 500 μL sample of undiluted plasma was spun at 1500 *x g* through the membrane and the chip was extracted from the device. The chip was allowed to dry and was then imaged by scanning electron microscopy as described below.

#### Tangential Flow for Analyte Capture

Nanoporous silicon nitride microfluidic devices were fabricated as described above. The NPN chip used had a 50 nm thick freestanding membrane with a 50 nm average pore diameter and a 15% porosity. 1 mL of plasma was passed tangential to the membrane surface at a rate of 10 μL/min using a syringe pump (Chemyx Fusion 200, Chemyx Inc., Stafford, TX), while fluid was actively pulled through the membrane at a rate of 2 μL/min. After processing the full 1 mL volume, the device was unclamped and the chip extracted. Captured sEVs were labeled for CD63 (Abcam, Cambridge, MA) and imaged via scanning electron microscopy as outlined below.

### Capture and Release

#### Microscale Experiments

Flow experiments were performed using two Chemyx Fusion 200 syringe pumps (Chemyx Inc., Stafford, TX). Micron scale experiments with 10 μm polystyrene green fluorescent particles (Thermo Scientific, USA) were conducted on 8 μm track-etch membranes. Capturing step was performed using a sample supply flow rate of 90 μL/min and an ultrafiltration/pulling rate of 10 μL/min. Captured particles were released by reversed flow of 10 μL/min through the membrane.

#### Nanoscale Experiments

These experiments were conducted using 100 nm polystyrene green fluorescent particles (Thermo Scientific, USA) on PCTE or NPN membranes with 80 nm median pore size. Nanoparticles were captured by supply flow rate of 5 μL/min and the ultrafiltration/pulling flow rate of 2 μL/min. Input channel was then cleaned by rinsing buffer to wash away the floating particles under the same flow condition as the capturing step. Finally, captured particles were released by reversed flow of 2 μL/min through the membranes.

### Time-Lapse Video Microscopy

Devices were illuminated with metal halide lamp source (LE6000 Leica) through DIC and FITC (488 nm Ex/525 nm Em) filter sets on a Leica DM16000 microscope (Leica Microsystems, Buffalo Grove, IL) using the 10X objective. Images were collected using MetaMorph software with a Rolera em-camera (QImaging, Surrey, BC Canada) for 50 ms exposure time for FITC and 10 ms for DIC. The measuring and merging channel tool in NIH ImageJ were used for quantifying the average intensity values and making videos by merging DIC with FITC images, respectively. Images were taken every minute for nanoscale experiments and every second for microscale experiments.

### Electron Microscopy

After the completion of experiments, the PCTE and NPN membranes were imaged via electron microscopy. Samples were prepared for electron microscopy by first removing the membranes from the device and then allowing them to air dry. Samples were then mounted and sputter coated with ~3-10 nm of gold. Scanning electron micrographs were taken at an accelerating voltage of 10 kV using either a Hitachi S-4000 scanning electron microscope (SEM) or a Zeiss AURIGA scanning electron microscope.

## Results

### Tangential Flow for Particle Capture

The system and scheme for particle capture and release is shown in Figure 1. As in our prior work (Burgin, Johnson, Chung, Clark, & McGrath, 2015; Chung et al., 2014; Mossu et al., 2018; Salminen et al., 2019), we used layer-by-layer assembly (Figure 1A) to construct microfluidic devices (Figure 1B) with membranes separating top and bottom flow channels. The only difference is that we used a clamped system for both PCTE systems and NPN systems instead of a fully bonded devices. This enables the removal and inspection of PCTE membranes or NPN chips by SEM after use. Particle capture (Figure 1C) was performed using two syringe pumps: a positive pressure pump providing a constant sample supply flow rate into the input channel of the device, and a negative pressure pump at the output channel exit side controlling a smaller, steady rate of ultrafiltration through the membrane. The difference between the supply and ultrafiltration rates exited the top channel as waste and provided the tangential flow needed to prevent fouling (Belfort et al., 1994; Field et al., 1995). The inlet port on the bottom channel was blocked for all experiments. After capture, non-adsorbed contaminants could be cleared by replacing the sample with a rinse buffer while maintaining the transmembrane pressure (Figure 1D) and captured analytes could be released by operating both pumps under positive pressure (Figure 1E).

### Small EV capture from Undiluted Serum

The initial discovery of analyte capture occurred with experiments on undiluted serum. In these experiments, we showed that the filtration of undiluted serum is difficult in NFF (Figure 2B), causing an 8 μm cake of serum protein and salts to foul the membrane and allowing the passage of only 10 μL of a 1 mL sample.

**Figure 2:**
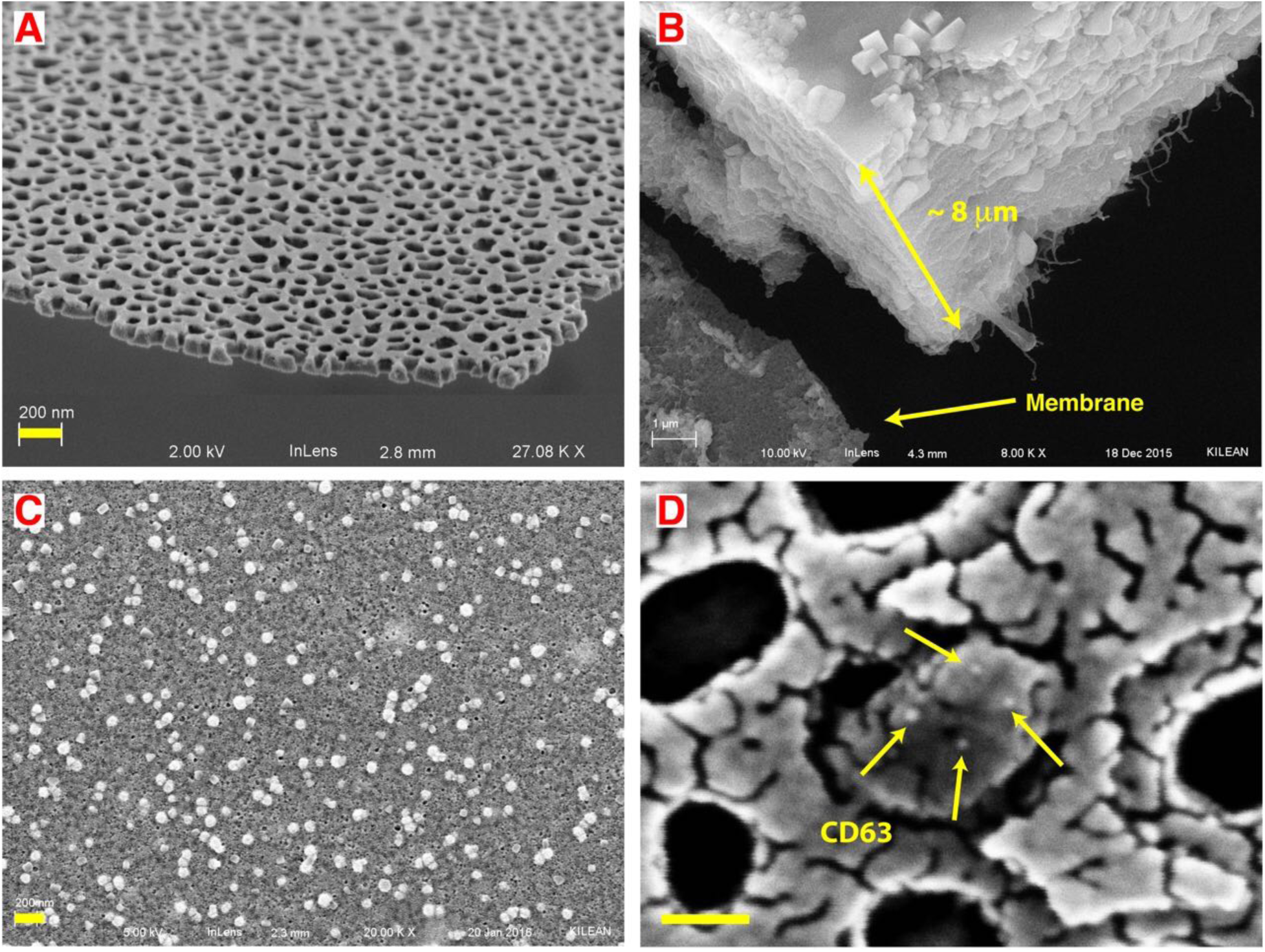
Small extracellular vesicles (sEV) captured from undiluted blood plasma. **A)** SEM images showing the thinness and high porosity of nanoporous silicon nitride (NPN). **B)** In normal flow filtration (NFF) a protein cake of ~8 μm cake rapidly builds up on the membrane surface. **C)** In tangential flow filtration (TFF) with the same sample, small vesicles are captured on the membrane surface with minimum fouling. **D)** Nanogold conjugated anti-CD63 antibody labels an EV captured in a pore multiple times, indicating it is likely a CD63 positive sEV. Note: the fragmented appearance of the surface results from the use of a limited amount of gold (3 nm) to avoid obscuring the gold label on the antibody (18 nm). By contrast 10 nm of gold was sputtered on the samples to avoid charging effects in SEM in both B and C. Scale bar = 200 nm for A, B and C. Scale bar = 50 nm for D.

However, upon passing the undiluted serum across the membrane in tangential flow, we observed a significant reduction in the protein build-up on the membrane, showing captured vesicles (Figure 2C). Further analysis of the vesicles with immunostaining showed that the vesicles were positive for CD63, a common sEV surface protein. We did not attempt rinse or release steps in with undiluted serum, instead we turned to the following experiments with model systems to confirm and study the capture phenomena under defined conditions.

### Microporous Track-Etch Capture of Fluorescent Particles

We first explored the particle capture phenomena at the microscale using microporous polycarbonate track-etched (mPCTE) membranes with 8 μm pores and 10 μm particles. At this scale we were able to image individual particle capture events in fluorescence microscopy (Figure 3). Before flow (T_0_; Figure 3B) there were no particles on the membrane. With a steady supply rate of 90 μL/min and ultrafiltration rate of 10 μL/min particles began to accumulate on the membrane, primarily drawn directly to the pores (see electron micrograph in Figure 3C, bottom panel), and the fluorescence steadily increased over time (Figure 3C). The capturing process was then stopped at T_1_ resulting in immediate release of particles loosely held on the membrane and a distinct, sudden drop in fluorescence (light blue line Figure 3A). Finally, the flow was reversed by switching the ultrafiltration pump to infusion mode, resulting in a directional shift for the bottom flow. The bottom flow rate was then increased to provide a high transmembrane pressure in an attempt to fully release the remaining particles, although a fraction remained irreversibly bound resulting in a residual fluorescence after the experiment T_2_(Figure 3D). Electron microscopy (Figure 3D, **bottom panel**) shows that most of these particles were not associated with pores and thus were non-specifically adhered to the surface of the membrane through surface interactions. More than 90% of the particles captured were released (**Supplementary Figure S2 and Movie S2**), suggesting this method has promise for the purification of microscale particles.

**Figure 3:**
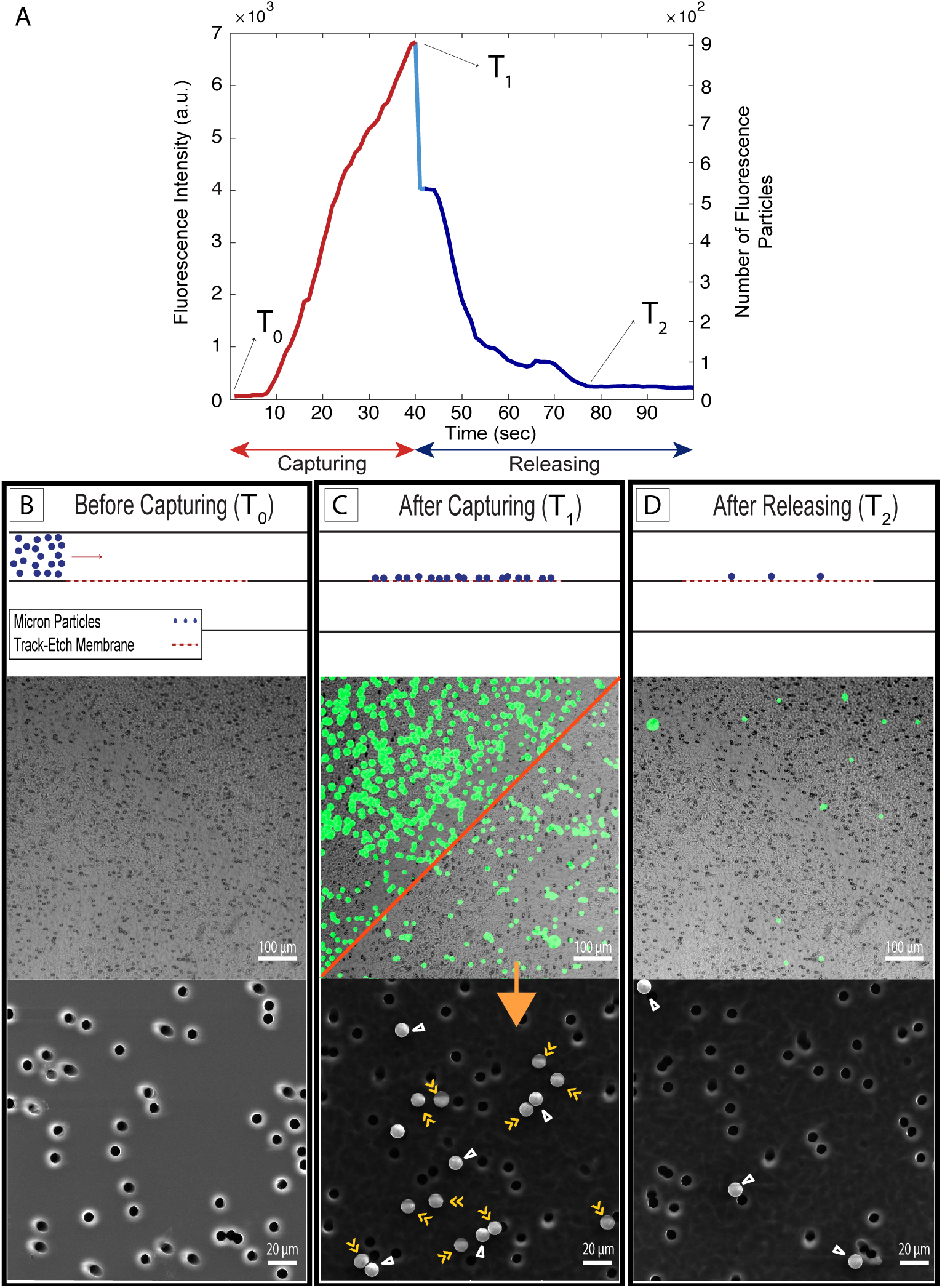
Microscale experiments with 10 μm fluorescent particles and 8 μm pore size polycarbonate track-etch (mPCTE) membranes. **A)** Fluorescence intensity/number of particles - time plot showing an increase in the intensity signal/number of particles during the capturing step and a decrease for releasing step. **B,C and D)** Before capturing, after capturing and after releasing panels, respectively, including schematic images (top), fluorescent images (middle) and scanning electron microscopy images (bottom). The red diagonal in fluorescent panel C shows releasing of captured particles by pausing the pump to change the flow configuration. Particles in panel C are labeled with either single or double arrowheads, indicating non-specifically bound and captured particles, respectively.

### Nanoporous Track-Etch Capture of Fluorescent Nanoparticles

Having demonstrated capture using modified tangential flow in a microscale system, experiments were performed to show capture and release at the nanoscale. Track-etch membranes with 80 nm pores (nPCTE) were used to capture 100 nm fluorescent nanoparticles. Because of the significant increase in membrane resistance compared to mPCTE, flow rates of 5 μL/min (sample supply) and 2 μL/min (ultrafiltration) were now used for capture. This was followed by washing with clean buffer to remove any non-specifically bound particles before the releasing in a backwash step (5 μL/min backflow).

During the capture phase of the experiments, the fluorescence intensity curves displayed similar behavior to the microscale experiments, with a steady increase throughout the capture period (Figure 4A). Unlike the mPCTE experiments however, there was no observable loss of fluorescence after the release of transmembrane pressure at the end of the capture phase. A fraction of loosely-associated particles, either on the surface or in suspension above the surface (Figure 4C), were removed with a wash step. Flow reversal did not fully remove all the particles captured on the membrane as some were lodged deep within pores (Figure 4D), but the system did return to within ~85% of the baseline fluorescence value.

**Figure 4:**
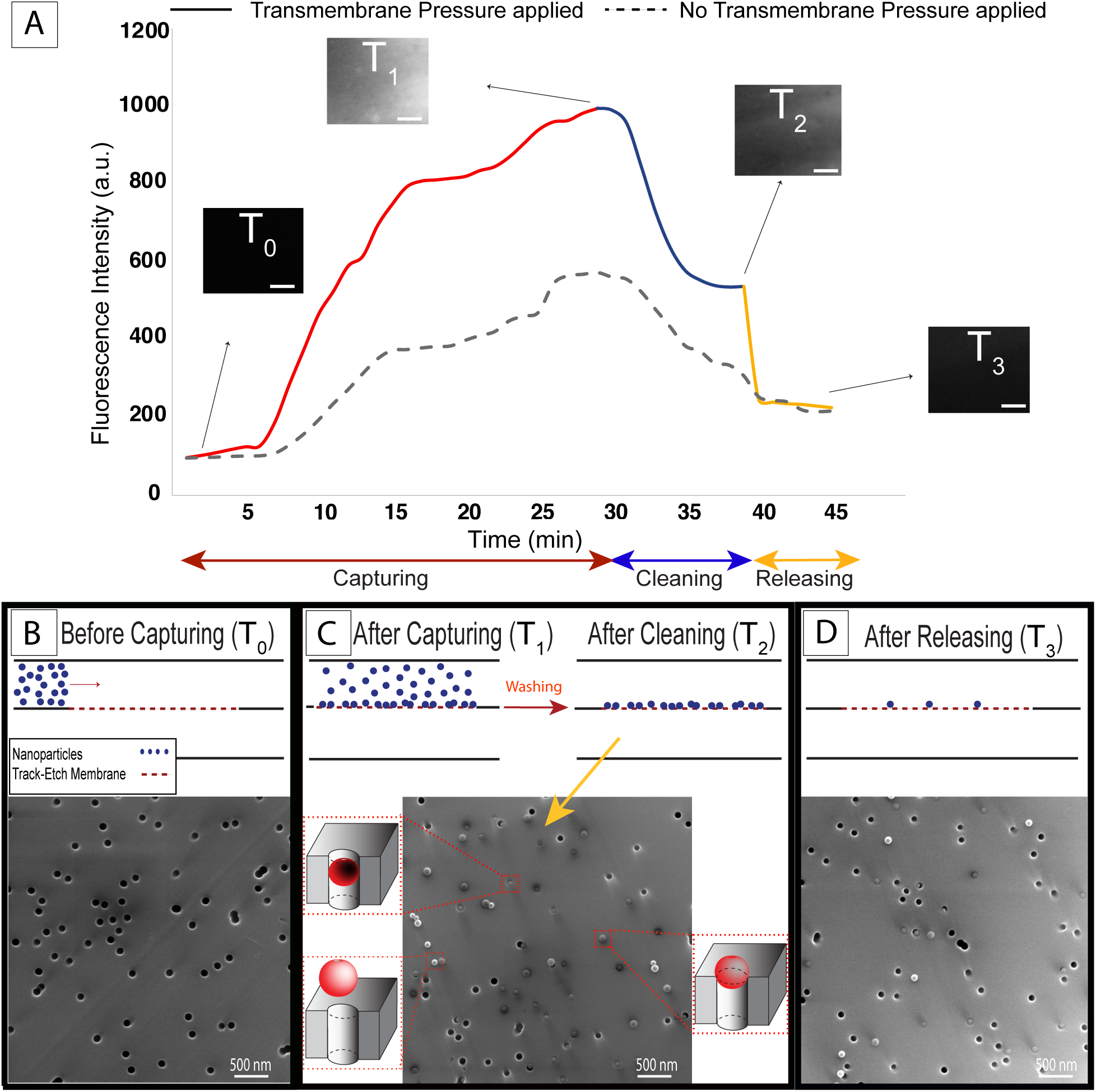
Nanoscale experiments with 100 nm fluorescent particles and 80 nm median pore size polycarbonate track-etch (nPCTE) membranes. **A)** Fluorescent intensity analysis (solid line) showing the gradual increasing and decreasing in the fluorescent signal during the capturing step and cleaning step, respectively, followed by a sharp drop as nanoparticles were released (the dash line shows the intensity change during the experiment in the absence of the transmembrane pressure). Scale bar on fluorescence image insets = 50 μm. **B, C and D)** Electron micrographs showing before capturing, after capturing-cleaning, and after releasing panels, respectively.

In order to assess the role of the applied transmembrane pressure on capturing, experiments were performed in the absence of active transmembrane pressure (dashed line, Figure 4A). To achieve this, supply flow was performed as before, but the ultrafiltration pump was not used to generate active transmembrane flow. While the change in fluorescence intensity showed an increase in particles, the maximum measured intensity was only 50% of the system with active transmembrane pressure which indicates that transmembrane pressure is the driving force of particle capture.

### Nanoporous Silicon Nanomembrane Capture of Fluorescent Nanoparticles

Our original observations of EV capture from serum (Figure 2) were obtained with 100 nm thick nanoporous silicon-nitride (NPN) membranes (J. P. S. DesOrmeaux et al., 2014). It is important to note that PCTE membranes used are approximately 60 times thicker compared to ultrathin nanomembranes. Thus, our next set of studies replicated the experimental conditions used with nPCTE on NPN (5 μL/min supply; 2 μL/min ultrafiltration) with similar pore sizes (80 nm median) and total number of pores actively filtering materials were of the same order (nPCTE = 4 × 10^7^ pores/mm^2^; NPN = 9.2 × 10^7^ pores/mm^2^), which resulted in a slightly larger membrane area for the nPCTE membranes (4 mm^2^) compared to the NPN membranes (1.4 mm^2^). Therefore, membrane thickness and membrane surface chemistry are the key parametric differences between experiments on nPCTE vs. NPN.

The capture and release intensity curves (Figure 5A) with NPN show similar trends to nPCTE with some interesting differences. There is again an increase in fluorescence intensity on the membrane during the capture phase followed by a sudden loss of particles when the flows are stopped. After a rinse with clean buffer, the intensity returns to within ~95% of the baseline, which is slightly better than that seen with nPCTE (Figure 5A, inset). A control in the absence of transmembrane pressure (Figure 5A, dashed line) showed once more that the capture process is driven by transmembrane pressure.

**Figure 5:**
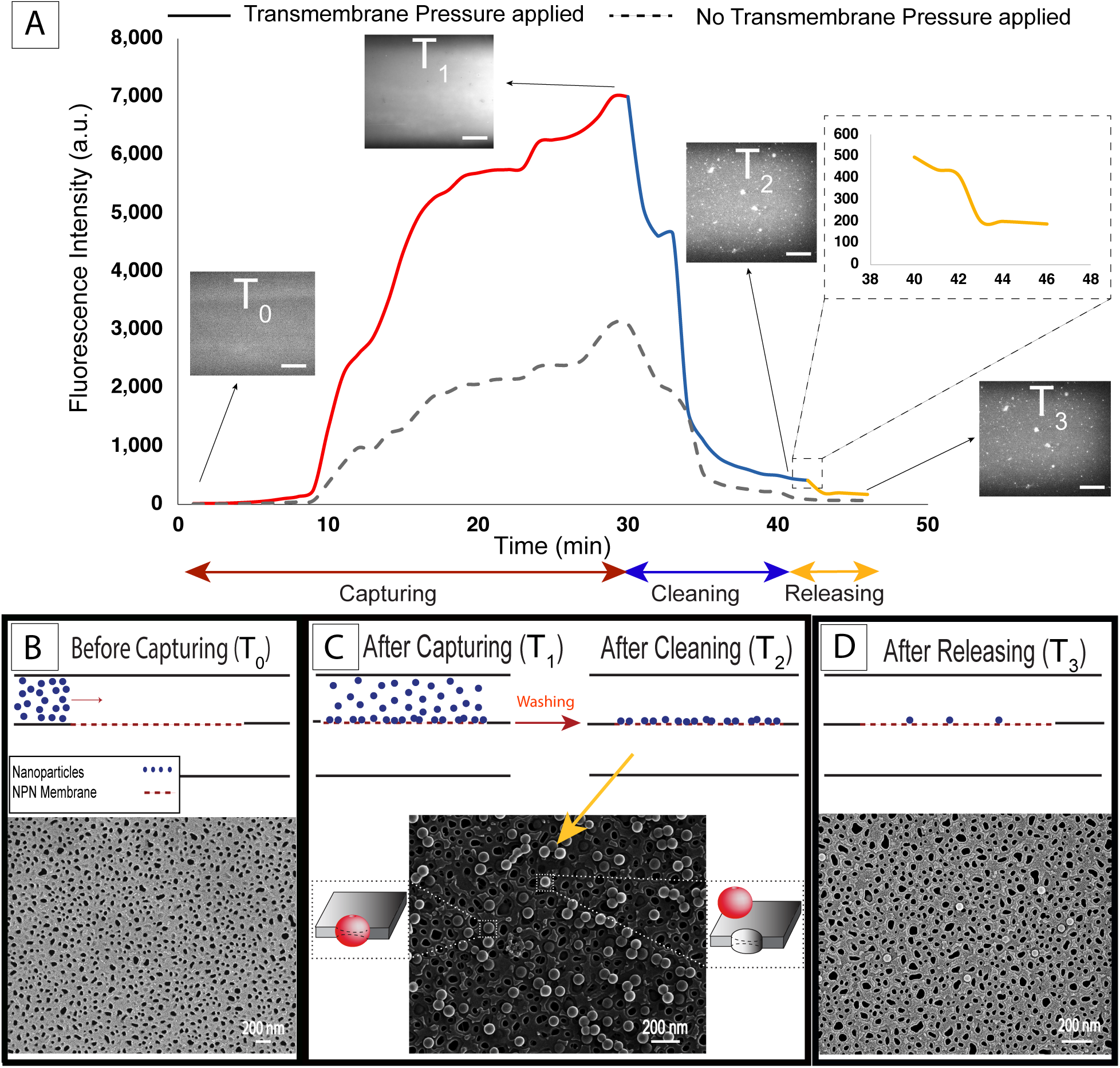
Nanoscale experiments with 100 nm diameter fluorescent particles and 80 nm median pore size nanoporous silicon nitride (NPN) membranes. **A)** Fluorescent intensity analysis (solid line) showing the increasing during the capturing step, and then decreasing in the cleaning step following by a drop as nanoparticles were released (the dash line showing the intensity changes during the experiment in the absence of the transmembrane pressure). Scale bar on fluorescence image insets = 50 μm. **B, C and D)** Electron micrographs showing before capturing, after capturing-cleaning, and after releasing panels respectively.

Electron microscopy was again performed to better understand the capture process. The membrane showed high pore density (Figure 5B), in contrast with track-etch membranes (Figure 4B), and a distribution of pore sizes with median of 80 nm (**Supplementary Figure S3**). As expected, the majority of the 100 nm particles captured remained on top of the pores (Figure 5C). A small proportion of particles persisted on the membrane after the releasing step, and these all appeared to be captured within pores (Figure 5D).

In order to estimate particle concentrations throughout the capture and release process, calibration curves for both NPN and nPCTE experiments were made by correlating the fluorescent intensity to the number of particles on the membrane (**Supplementary Figure S4**). These curves allowed for the direct comparison of membrane performance for particle capture and release (Table 1). We estimate that track-etch membranes capture ~2.6 × 10^6^ particles from an available population of 5 × 10^7^ and released 60% of the particles captured. By contrast, silicon nanomembranes captured ~8.6 × 10^6^ particles from the same solution and released 68% of the captured population.

**Table 1:** Nanoporous track-etch (nPCTE) vs. silicon nanomembrane (NPN) captured and released particle counts.

| Membrane | Captured Particles | Released Particles |
| --- | --- | --- |
| TE Membranes | $(2.58 \pm 0.24) \times 10^6$ | $(1.56 \pm 0.43) \times 10^6$ |
| NPN Membranes | $(8.64 \pm 3.15) \times 10^6$ | $(5.89 \pm 1.98) \times 10^6$ |

### Pressure Modeling of Track-Etch and Ultrathin Silicon Nitride Nanomembranes

We explored the effect of the membrane thickness on transmembrane pressure in our studies both analytically and experimentally. Experiments were conducted with nanoporous track-etch membranes and ultrathin nanoporous silicon nitride nanomembranes. Pressure sensors were placed upstream and downstream on either side of the membrane (Figure 6A) and the pressures were monitored under flow conditions equivalent to the capture experiments (5 μL/min supply, 2 μL/min ultrafiltration). Results for both nPCTE and NPN compared favorably to predictions of the Dagan equation – a modified Hagen-Poiseuille equation that also applies to ultrathin membranes (Chung et al., 2014; Dagan, Weinbaum, & Pfeffer, 1983; Gaborski et al., 2010). The Dagan equation gives the pore resistance as:

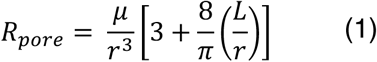

where *μ* is the fluid viscosity [Pa s^−1^], *r* is the pore radius [m], and *L* is the pore length [m]. The total membrane resistance *R* is calculated by adding the resistance for each pore in the membrane in parallel (9.2 × 10^7^ pores for NPN and 4 × 10^7^ pores for nPCTE) and the anticipated pressure drop is then found by multiplying by the flow rate:

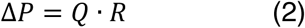

**Figure 6:**
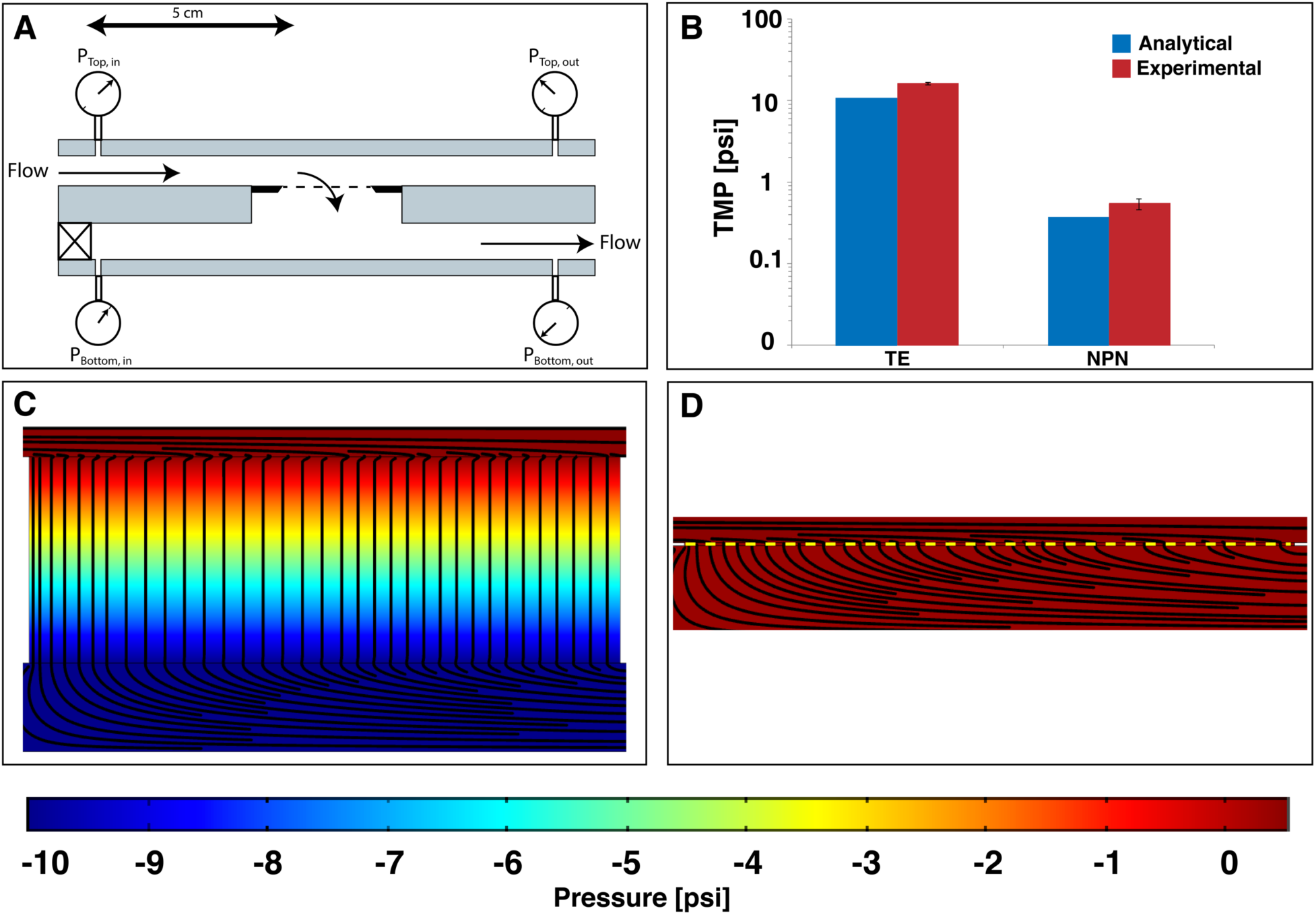
Theoretical and experimental pressure drops across nanoporous polycarbonate track-etch membranes (nPCTE) and nanoporous silicon nitride (NPN) membranes. **A)** Diagram of the pressure monitoring system showing the position of the pressure sensors and the direction of flow. All flow was performed at 10 μL/min through the membrane with a syringe pump pushing on the top channel and a syringe pump pulling on the bottom channel. The pressure sensors were positioned 5 cm above and below the membrane. **B)** Comparison of pressure drops across the track-etch and NPN membranes. Blue = Dagan predicted, homogeneous distribution pressure drop. Red = experimental data. Logarithmic scale used for comparison. **C)** COMSOL model of pressure in a track-etch system showing a large pressure drop across the membrane. **D)** COMSOL model of pressure in an NPN system showing almost no pressure drop across the membrane, in stark contrast to the track etch system. COMSOL simulations were performed using the Free and Porous Flow toolbox with a Darcy’s permeability calculated for this system.

The comparison of this estimate with experimental results (Figure 6B) showed that a simple analytical approximation is sufficient for predicting the transmembrane pressure drop that could be experienced in the system. These results were compared to an analytical model of pressure drop (Figure 6B) as well as COMSOL Multiphysics simulations (Figure 6C and **6D**) to illustrate the pressure gradients and streamlines in the system.

## Discussion

In this work, we introduced a new method for sample purification in which particles are captured on the surface of a membrane in tangential flow, washed to remove contaminants, and then released in a controlled fashion where they can be further analyzed, concentrated or processed.

We call this process tangential flow for analyte capture (TFAC) and while the process resembles bind and elute strategies found in column or membrane chromatography, it relies on physical interaction, rather than chemical affinity, for capture. Similarly, TFAC requires physical release through back-flow for elution, rather than chemically treatments to disassociate chemical bonds formed during capture. As the release of chemical bonds in affinity schemes can often be destructive and incomplete, there are clear advantages for physical capture and release.

Proof of concept experiments using fluorescent particles on both PCTE and NPN membranes showed successful capture and release of particles. We have shown that NPN membranes outperform PCTE membranes for capture and release with polystyrene nanoparticles. Our analytical and experimental comparison showed that the greater thickness of PCTE compared to NPN caused higher transmembrane pressure. This high pressure drives nanoparticles into the membrane bulk where they disappear from view and are more difficult to recover (Figure 4).

One potential application of TFAC utilizing ultrathin membranes as a microfluidic based technique would be isolation of extracellular vesicles. Currently, the “gold standard” method for isolating extracellular vesicles from biofluids is ultracentrifugation, which requires large volumes of biofluid (> 25 ml), long processing times, expensive instrumentation and trained technicians. Gel precipitation and size exclusion chromatography and have been developed that remove the need for ultracentrifugation and allow extracellular vesicle isolation in a benchtop centrifuge (Baranyai et al., 2015; Böing et al., 2014; Enderle et al., 2015; Wu & Antes, 2010), but these methods suffer from low yield and/or contamination with co-precipitated proteins (Baranyai et al., 2015; Enderle et al., 2015; Lee et al.). The high protein contamination from these methods prevents the use of EV proteins as biomarkers in addition to RNA. The result from our plasma isolation experiment by ultrathin nanomembranes showing capture of EVs with minimal contamination suggests promising potential of TFAC for isolation of EVs with high purity (Figure 7).

**Figure 7:**
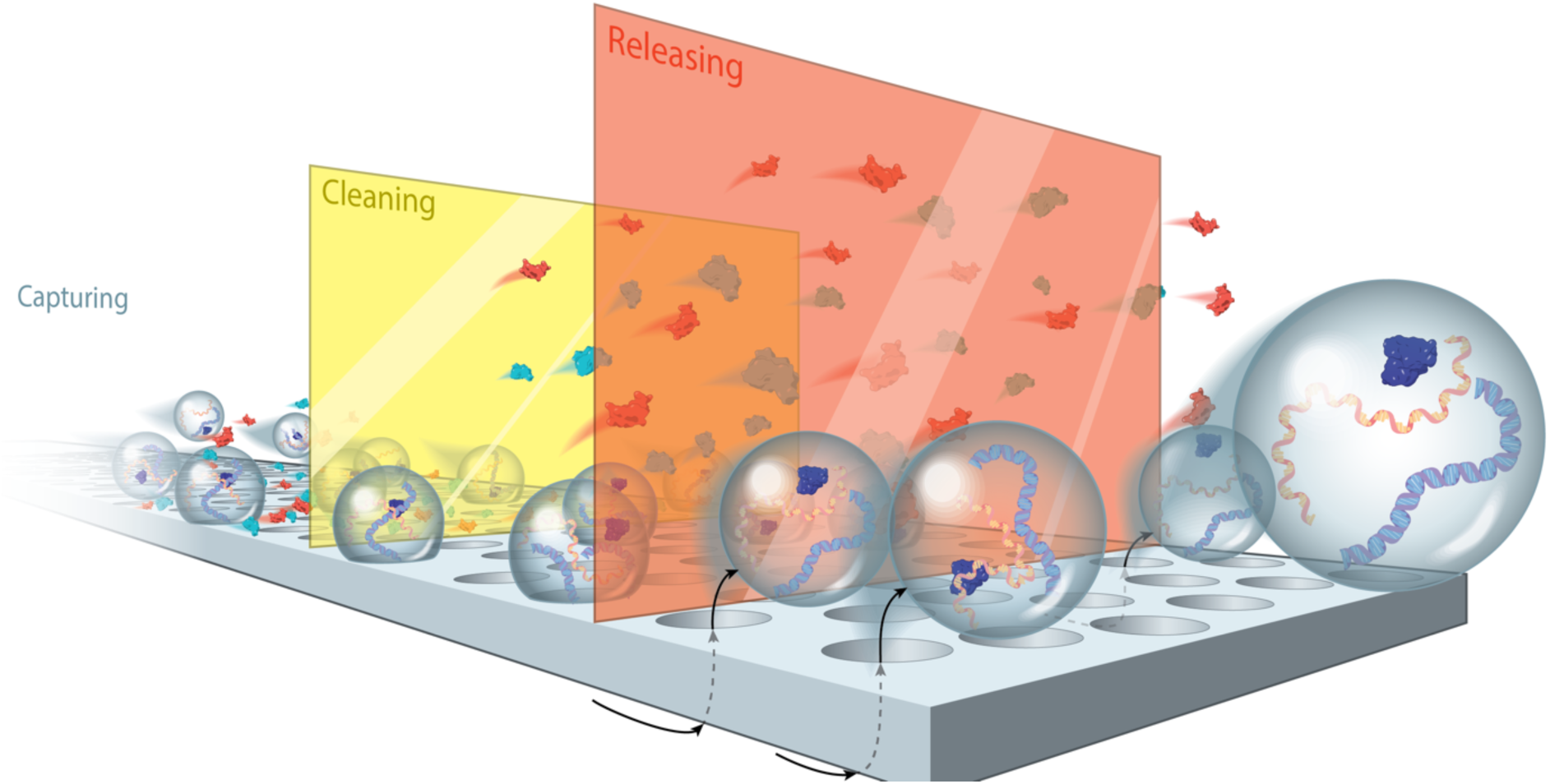
Tangential Flow for Analyte Capture (TFAC) Illustration showing capturing, cleaning and releasing steps.

Additionally, the pore size of the ultrathin NPN membranes can be tuned to capture different subpopulations of EVs that vary in size (Fang et al., 2010, Jamaly et al., 2018). This includes microvesicles, exosomes, apoptotic bodies and lipoprotein particles which are diagnostically informative (Clayton et al., 2018). In all cases, TFAC method eliminates the necessity of preprocessing biofluids which can be both time consuming, result in sample loss, and often requires specialized equipment reducing the utility of these particles in point of care diagnostic devices (Soekmadji et al., 2018, Das et al., 2019).

Another potential application of TFAC would be a membrane-based ‘*in situ’* analysis to detect EVs carrying cancer biomarkers among a larger population using the same membrane for capture, labeling, and imaging by fluorescence microscopy. TFAC using NPN membranes showed that captured particles were associated with membrane surface, rather than trapped in a bulk-matrix which means that the captured particles can be analyzed directly on the membrane. Furthermore, TFAC captured extracellular vesicles from whole plasma with minimal contamination (Figure 2C) as opposed to rapidly formed cake on the membrane by NFF which increases the sensitivity and specify of EVs biomarker detection. Also, the excellent optical properties of ultrathin inorganic membranes like NPN (Carter et al., 2017), would also be key to enabling this application. In comparison, track-etch membranes lack this optical transparency and as the current study indicates, trap EVs below the membrane surface, together precluding the ability to detect specific diagnostic markers directly in and on the EVs captured on the membranes (Martínez-Pérez and García-Rupérez, 2019).

## Conclusion

In this work, we have developed a method called tangential flow for analyte capture (TFAC) to capture and release of particles. We contend that ultrathin membranes are ideally suited for TFAC for two reasons: 1) operating pressures are orders-of-magnitude lower for ultrathin membranes than for membranes with conventional thicknesses (1-10 μm) and 2) captured particles are associated with a surface, rather than trapped in a bulk-matrix and 3) higher efficiency of capture and release of particles. Experiments performed in normal flow filtration with human plasma demonstrated formation of a protein cake on the surface of ultrathin membranes. However, testing human plasma in TFAC mode resulted in capturing extracellular vesicles with minimal contamination. Captured vesicles were further labeled *in situ*, providing a convenient platform for downstream detection and analysis. Together, these findings suggest promising potential of TFAC for both isolation of EVs and biomarker detection on captured EVs.

## Acknowledgements and Funding

The authors would like to acknowledge Brad Kwarta for the EV capture and release illustration. Research reported in this publication was supported in part by the National Science Foundation (IIP 1660177) to J.L.M and T.R.G., Department of Defense (CA170373) to J.L.M., and the National Institutes of Health (R35GM119623) to T.R.G.

## References

Armstrong, D., & Wildman, D. E. (2018). Extracellular Vesicles and the Promise of Continuous Liquid Biopsies. J Pathol Transl Med, 52(1), 1–8. doi: 10.4132/jptm.2017.05.21

Baranyai, T., Herczeg, K., Onódi, Z., Voszka, I., Módos, K., Marton, N., … Giricz, Z. (2015). Isolation of Exosomes from Blood Plasma: Qualitative and Quantitative Comparison of Ultracentrifugation and Size Exclusion Chromatography Methods. PLOS ONE, 10(12), e0145686. doi: 10.1371/journal.pone.0145686

Belfort, G., Davis, R. H., & Zydney, A. L. (1994). The behavior of suspensions and macromolecular solutions in crossflow microfiltration. J Memb Sci, 96, 1–58.

Böing, A. N., van der Pol, E., Grootemaat, A. E., Coumans, F. A. W., Sturk, A., & Nieuwland, R. (2014). Single-step isolation of extracellular vesicles by size-exclusion chromatography. Journal of Extracellular Vesicles, 3, 10.3402/jev.v3403.23430. doi: 10.3402/jev.v3.23430

Burgin, T., Johnson, D., Chung, H., Clark, A., & McGrath, J. (2015). Analytical and Finite Element Modeling of Nanomembranes for Miniaturized, Continuous Hemodialysis. Membranes, 6(1), 6.

Burkhart, C. T., Maki, K. L., & Schertzer, M. J. (2017). Effects of Interface Velocity, Diffusion Rate, and Radial Velocity on Colloidal Deposition Patterns Left by Evaporating Droplets. Journal of Heat Transfer, 139(11), 111505.

Busatto, S., Vilanilam, G., Ticer, T., Lin, W. L., Dickson, D. W., Shapiro, S., … Wolfram, J. (2018). Tangential Flow Filtration for Highly Efficient Concentration of Extracellular Vesicles from Large Volumes of Fluid. Cells, 7(12). doi: 10.3390/cells7120273

Carter, R. N., Casillo, S. M., Mazzocchi, A. R., DesOrmeaux, J. S., Roussie, J. A., & Gaborski, T. R. (2017). Ultrathin transparent membranes for cellular barrier and co-culture models. Biofabrication, 9(1), 015019. doi: 10.1088/1758-5090/aa5ba7

Christy, C., Adams, G., Kuriyel, R., Bolton, G., & Seilly, A. (2002). High-performance tangential flow filtration: a highly selective membrane separation process. Desalination, 144(1-3), 133–136.

Chung, H. H., Chan, C. K., Khire, T. S., Marsh, G. A., Clark, A., Waugh, R. E., & McGrath, J. L. (2014). Highly permeable silicon membranes for shear free chemotaxis and rapid cell labeling. Lab Chip, 14, 2456–2468. doi: 10.1039/c4lc00326h

Clayton, A., Buschmann, D., Byrd, J. B., Carter, D. R. F., Cheng, L., Compton, C., … Nieuwland, R. (2018). Summary of the ISEV workshop on extracellular vesicles as disease biomarkers, held in Birmingham, UK, during December 2017. J Extracell Vesicles, 7(1), 1473707.

Conlan, R. S., Pisano, S., Oliveira, M. I., Ferrari, M., & Pinto, I. M. (2017). Exosomes as reconfigurable therapeutic systems. Trends in molecular medicine, 23(7), 636–650.

Dagan, Z., Weinbaum, S., & Pfeffer, R. (1983). Theory and experiment on the three-dimensional motion of a freely suspended spherical particle at the entrance to a pore at low Reynolds number. Chemical Engineering Science, 38(4), 583–596. doi: https://doi.org/10.1016/0009-2509(83)80118-3

Das, S., Extracellular, R. N. A. C. C., Ansel, K. M., Bitzer, M., Breakefield, X. O., Charest, A., … Laurent, L. C. (2019). The Extracellular RNA Communication Consortium: Establishing Foundational Knowledge and Technologies for Extracellular RNA Research. Cell, 177(2), 231–242.

Davies, R. T., Kim, J., Jang, S. C., Choi, E.-J., Gho, Y. S., & Park, J. (2012). Microfluidic filtration system to isolate extracellular vesicles from blood. Lab on a Chip, 12(24), 5202–5210. doi: 10.1039/C2LC41006K

DesOrmeaux, J. P., Winans, J. D., Wayson, S. E., Gaborski, T. R., Khire, T. S., Striemer, C. C., & McGrath, J. L. (2014). Nanoporous silicon nitride membranes fabricated from porous nanocrystalline silicon templates. Nanoscale, 6(18), 10798–10805. doi: 10.1039/c4nr03070b

DesOrmeaux, J. P. S., Winans, J. D., Wayson, S. E., Gaborski, T. R., Khire, T. S., Striemer, C. C., & McGrath, J. L. (2014). Nanoporous silicon nitride membranes fabricated from porous nanocrystalline silicon templates. Nanoscale, 6(18), 10798–10805. doi: Doi 10.1039/C4nr03070b

Enderle, D., Spiel, A., Coticchia, C. M., Berghoff, E., Mueller, R., Schlumpberger, M., … Noerholm, M. (2015). Characterization of RNA from Exosomes and Other Extracellular Vesicles Isolated by a Novel Spin Column-Based Method. PLOS ONE, 10(8), e0136133. doi: 10.1371/journal.pone.0136133

Fang, D. Z., Striemer, C. C., Gaborski, T. R., McGrath, J. L., & Fauchet, P. M. (2010). Methods for controlling the pore properties of ultra-thin nanocrystalline silicon membranes. J Phys Condens Matter, 22(45), 454134

Fernando, M. R., Jiang, C., Krzyzanowski, G. D., & Ryan, W. L. (2017). New evidence that a large proportion of human blood plasma cell-free DNA is localized in exosomes. PLoS One, 12(8), e0183915. doi: 10.1371/journal.pone.0183915

Field, R. W., Wu, D., Howell, J. A., & Gupta, B. B. (1995). Critical Flux Concept for Microfiltration Fouling. Journal of Membrane Science, 100(3), 259–272.

Fritsch, J., & Moraru, C. I. (2008). Development and optimization of a carbon dioxide-aided cold microfiltration process for the physical removal of microorganisms and somatic cells from skim milk. Journal of Dairy Science, 91(10), 3744–3760.

Gaborski, T. R., Snyder, J. L., Striemer, C. C., Fang, D. Z., Hoffman, M., Fauchet, P. M., & McGrath, J. L. (2010). High-performance separation of nanoparticles with ultrathin porous nanocrystalline silicon membranes. ACS Nano, 4(11), 6973–6981. doi: 10.1021/nn102064c

Gallo, A., Tandon, M., Alevizos, I., & Illei, G. G. (2012). The majority of microRNAs detectable in serum and saliva is concentrated in exosomes. PLoS One, 7(3), e30679. doi: 10.1371/journal.pone.0030679

Ghosh, R. (2006). Rapid antibody screening by membrane chromotagraphic immunoassay technique. J Chromatogr B, 844, 163–167.

Harris, S. G., & Shuler, M. L. (2003). Growth of endothelial cells on microfabricated silicon nitride membranes for an in vitro model of the blood-brain barrier. Biotechnology and Bioprocess Engineering, 8(4), 246–251.

Huang, X., Yuan, T., Tschannen, M., Sun, Z., Jacob, H., Du, M., … Wang, L. (2013). Characterization of human plasma-derived exosomal RNAs by deep sequencing. BMC Genomics, 14(1), 319. doi: 10.1186/1471-2164-14-319

Jamaly, S., Ramberg, C., Olsen, R., Latysheva, N., Webster, P., Sovershaev, T., … Hansen, J. B. (2018). Impact of preanalytical conditions on plasma concentration and size distribution of extracellular vesicles using Nanoparticle Tracking Analysis. Sci Rep, 8(1), 17216.

Johnson, D. G., Khire, T. S., Lyubarskaya, Y. L., Smith, K. J., Desormeaux, J. P., Taylor, J. G., … McGrath, J. L. (2013). Ultrathin silicon membranes for wearable dialysis. Adv Chronic Kidney Dis, 20(6), 508–515. doi: 10.1053/j.ackd.2013.08.001

Kang, D., Oh, S., Ahn, S. M., Lee, B. H., & Moon, M. H. (2008). Proteomic analysis of exosomes from human neural stem cells by flow field-flow fractionation and nanoflow liquid chromatography-tandem mass spectrometry. J Proteome Res, 7(8), 3475–3480. doi: 10.1021/pr800225z

Kurnik, R. T., Yu, A. W., Blank, G. S., Burton, A. R., Smith, D., Athalye, A. M., & van Reis, R. (1995). Buffer exchange using size exclusion chromatography, countercurrent dialysis, and tangential flow filtration: Models, development, and industrial application. Biotechnol Bioeng, 45(2), 149–157. doi: 10.1002/bit.260450209

Lane, R. E., Korbie, D., Hill, M. M., & Trau, M. (2018). Extracellular vesicles as circulating cancer biomarkers: opportunities and challenges. Clin Transl Med, 7(1), 14. doi: 10.1186/s40169-018-0192-7

Langfield, K. K., Walker, H. J., Gregory, L. C., & Federspiel, M. J. (2011). Manufacture of Measles Viruses. In O.-W. Merten & M. Al-Rubeai (Eds.), Viral Vectors for Gene Therapy: Methods and Protocols (Vol. 737, pp. 345–366): Humana Press.

Lee, C., Carney Rp Fau - Hazari, S., Hazari S Fau - Smith, Z. J., Smith Zj Fau - Knudson, A., Knudson A Fau - Robertson, C. S., Robertson Cs Fau - Lam, K. S., … Wachsmann-Hogiu, S. 3D plasmonic nanobowl platform for the study of exosomes in solution. (2040–3372 (Electronic)).

Li, M., Zeringer, E., Barta, T., Schageman, J., Cheng, A., & Vlassov, A. V. (2014). Analysis of the RNA content of the exosomes derived from blood serum and urine and its potential as biomarkers. Philos Trans R Soc Lond B Biol Sci, 369(1652). doi: 10.1098/rstb.2013.0502

Li, P., Kaslan, M., Lee, S. H., Yao, J., & Gao, Z. (2017). Progress in Exosome Isolation Techniques. Theranostics, 7(3), 789–804. doi: 10.7150/thno.18133

Li, Y., Yang, Q., Li, M., & Song, Y. (2016). Rate-dependent interface capture beyond the coffee-ring effect. Scientific reports, 6, 24628.

Lock, M., Alvira, M. R., & Wilson, J. M. (2012). Analysis of particle content of recombinant adeno-associated virus serotype 8 vectors by ion-exchange chromatography. Hum Gene Ther Methods, 23(1), 56–64. doi: 10.1089/hgtb.2011.217

Martinez-Perez, P., & Garcia-Ruperez, J. (2019). Commercial polycarbonate track-etched membranes as substrates for low-cost optical sensors. Beilstein J Nanotechnol, 10, 677–683.F

Mathivanan, S., Lim, J. W. E., Tauro, B. J., Ji, H., Moritz, R. L., & Simpson, R. J. (2010). Proteomics Analysis of A33 Immunoaffinity-purified Exosomes Released from the Human Colon Tumor Cell Line LIM1215 Reveals a Tissue-specific Protein Signature. Molecular & Cellular Proteomics: MCP, 9(2), 197–208. doi: 10.1074/mcp.M900152-MCP200

Mazzocchi, A. R., Man, A. J., DesOrmeaux, J. P. S., & Gaborski, T. R. (2014). Porous Membranes Promote Endothelial Differentiation of Adipose-Derived Stem Cells and Perivascular Interactions. Cellular and Molecular Bioengineering, 7(3), 369–378.

McNamara, R. P., Caro-Vegas, C. P., Costantini, L. M., Landis, J. T., Griffith, J. D., Damania, B. A., & Dittmer, D. P. (2018). Large-scale, cross-flow based isolation of highly pure and endocytosis-competent extracellular vesicles. J Extracell Vesicles, 7(1), 1541396. doi: 10.1080/20013078.2018.1541396

Mireles, M., & Gaborski, T. R. (2017). Fabrication techniques enabling ultrathin nanostructured membranes for separations. Electrophoresis, 38(19), 2374–2388.

Mossu, A., Rosito, M., Khire, T., Li Chung, H., Nishihara, H., Gruber, I., … Engelhardt, B. (2018). A silicon nanomembrane platform for the visualization of immune cell trafficking across the human blood-brain barrier under flow. J Cereb Blood Flow Metab, 271678X18820584. doi: 10.1177/0271678X18820584

Nair, R. R., Wu, H. A., Jayaram, P. N., Grigorieva, I. V., & Geim, A. K. (2012). Unimpeded permeation of water through helium-leak-tight graphene-based membranes. Science, 335(6067), 442–444. doi: 10.1126/science.1211694

Okada, T., Nonaka-Sarukawa, M., Uchibori, R., Kinoshita, K., Hayashita-Kinoh, H., Nitahara-Kasahara, Y., … Ozawa, K. (2009). Scalable purification of adeno-associated virus serotype 1 (AAV1) and AAV8 vectors, using dual ion-exchange adsorptive membranes. Hum Gene Ther, 20(9), 1013–1021. doi: 10.1089/hum.2009.006

Rekker, K., Saare, M., Roost, A. M., Kubo, A.-L., Zarovni, N., Chiesi, A., … Peters, M. (2014). Comparison of serum exosome isolation methods for microRNA profiling. Clinical Biochemistry, 47(1), 135–138. doi: https://doi.org/10.1016/j.clinbiochem.2013.10.020

Salminen, A. T., Zhang, J., Madejski, G. R., Khire, T. S., Waugh, R. E., McGrath, J. L., & Gaborski, T. R. (2019). Ultrathin Dual-Scale Nano- and Microporous Membranes for Vascular Transmigration Models. Small, e1804111. doi: 10.1002/smll.201804111

Segura, M., Kamen, A., & Garnier, A. (2011). Overview of Current Scalable Methods for Purification of Viral Vectors. In O.-W. Merten & M. Al-Rubeai (Eds.), Viral Vectors for Gene Therapy: Methods and Protocols (Vol. 737, pp. 89–116): Humana Press.

Smith, K. J., Winans, J., & McGrath, J. (2016). Ultrathin Membrane Fouling Mechanism Transitions in Dead-End Filtration of Protein. Paper presented at the ASME 2016 14th International Conference on Nanochannels, Microchannels, and Minichannels collocated with the ASME 2016 Heat Transfer Summer Conference and the ASME 2016 Fluids Engineering Division Summer Meeting.

Smith, K. J. P., May, M., Baltus, R. E., & McGrath, J. L. (2017). A predictive model of separations in dead-end filtration with ultrathin membranes. Separation and Purification Technology, 189, 40–47.

Snyder, J. L., Clark, A., Jr., Fang, D. Z., Gaborski, T. R., Striemer, C. C., Fauchet, P. M., & McGrath, J. L. (2011). An experimental and theoretical analysis of molecular separations by diffusion through ultrathin nanoporous membranes. J Memb Sci, 369(1-2), 119–129. doi: 10.1016/j.memsci.2010.11.056

Soekmadji, C., Hill, A. F., Wauben, M. H., Buzas, E. I., Di Vizio, D., Gardiner, C., … Witwer, K. W. (2018). Towards mechanisms and standardization in extracellular vesicle and extracellular RNA studies: results of a worldwide survey. J Extracell Vesicles, 7(1), 1535745.

Striemer, C. C., Gaborski, T. R., McGrath, J. L., & Fauchet, P. M. (2007). Charge- and size-based separation of macromolecules using ultrathin silicon membranes. Nature, 445(7129), 749–753. doi: Doi 10.1038/Nature05532

Surwade, S. P., Smirnov, S. N., Vlassiouk, I. V., Unocic, R. R., Veith, G. M., Dai, S., & Mahurin, S. M. (2015). Water desalination using nanoporous single-layer graphene. Nat Nanotechnol, 10(5), 459–464. doi: 10.1038/nnano.2015.37

Sweeny, S., Woehrle, G., & Hutchinson, J. (2006). Rapid Purification and Size Separation of Gold Nanoparticles via Diafiltration. Journal of American Chemical Society, 128, 3190–3197.

Taylor, D. D., & Gercel-Taylor, C. (2008). MicroRNA signatures of tumor-derived exosomes as diagnostic biomarkers of ovarian cancer. Gynecologic Oncology, 110(1), 13–21. doi: https://doi.org/10.1016/j.ygyno.2008.04.033

Théry, C., Amigorena, S., Raposo, G., & Clayton, A. (2001). Isolation and Characterization of Exosomes from Cell Culture Supernatants and Biological Fluids Current Protocols in Cell Biology: John Wiley & Sons, Inc.

Théry, C., Witwer, K. W., Aikawa, E., Alcaraz, M. J., Anderson, J. D., Andriantsitohaina, R., … Zuba-Surma, E. K. (2018). Minimal information for studies of extracellular vesicles 2018 (MISEV2018): a position statement of the International Society for Extracellular Vesicles and update of the MISEV2014 guidelines. Journal of extracellular vesicles, 7(1), 1535750. doi: 10.1080/20013078.2018.1535750

van Reis, R., Gadam, S., Frautschy, L. N., Orlando, S., Goodrich, E. M., Saksena, S., … Zydney, A. L. (1997). High performance tangential flow filtration. Biotechnol Bioeng, 56(1), 71–82. doi: 10.1002/(SICI)1097-0290(19971005)56:1<71::AID-BIT8>3.0.CO;2-S

van Reis, R., & Zydney, A. (2001). Membrane separations in biotechnology. Current Opinion in Biotechnology, 12(2), 208–211. doi: Doi 10.1016/S0958-1669(00)00201-9

Vlassiouk, I., Apel, P. Y., Dmitriev, S. N., Healy, K., & Siwy, Z. S. (2009). Versatile ultrathin nanoporous silicon nitride membranes. Proc Natl Acad Sci U S A, 106(50), 21039–21044. doi: 10.1073/pnas.09114501060911450106[pii]

Winans, J., Smith, K., Gaborski, T., Roussie, J., & McGrath, J. (2016). Membrane capacity and fouling mechanisms for ultrathin nanomembranes in dead-end filtration. J Memb Sci, 499, 282–289.

Wu, F., & Antes, T. J. (2010). Abstract 3030: An exosome isolation system for serum-based cancer biomarker discovery. Cancer Research, 70(8 Supplement), 3030.

Yuen, P. K., & Goral, V. N. (2010). Low-cost rapid prototyping of flexible microfluidic devices using a desktop digital craft cutter. Lab on a Chip, 10(3), 384–387.

