## Supplemental Figures for "Tangential flow microfluidics for the capture and release of nanoparticles and extracellular vesicles on conventional and ultrathin membranes"

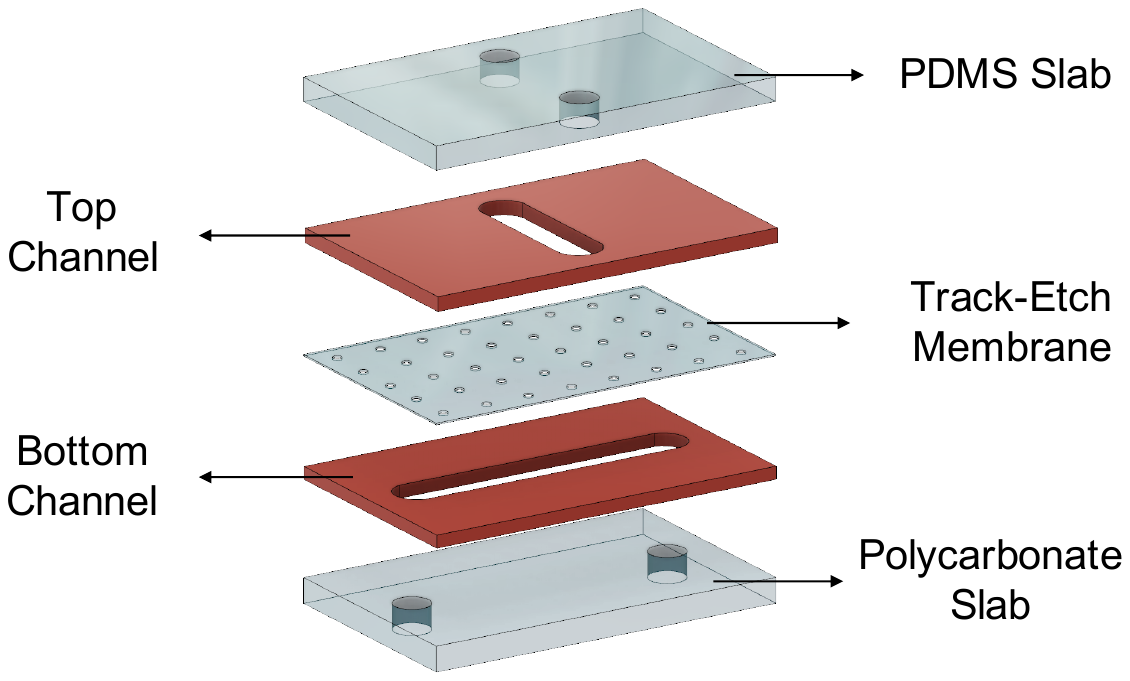

**Supplementary Figure 1; Microfluidic device for PCTE membranes**. Polycarbonate and PDMS slabs with holes were used to have access to the bottom and top channels respectively. Top and bottom channels were patterned into PDMS sheets, and PCTE membrane was sandwiched between the channels. In order to make sure that the system is sealed, a plus sign design for channels were used and the PCTE was covering the entire device.

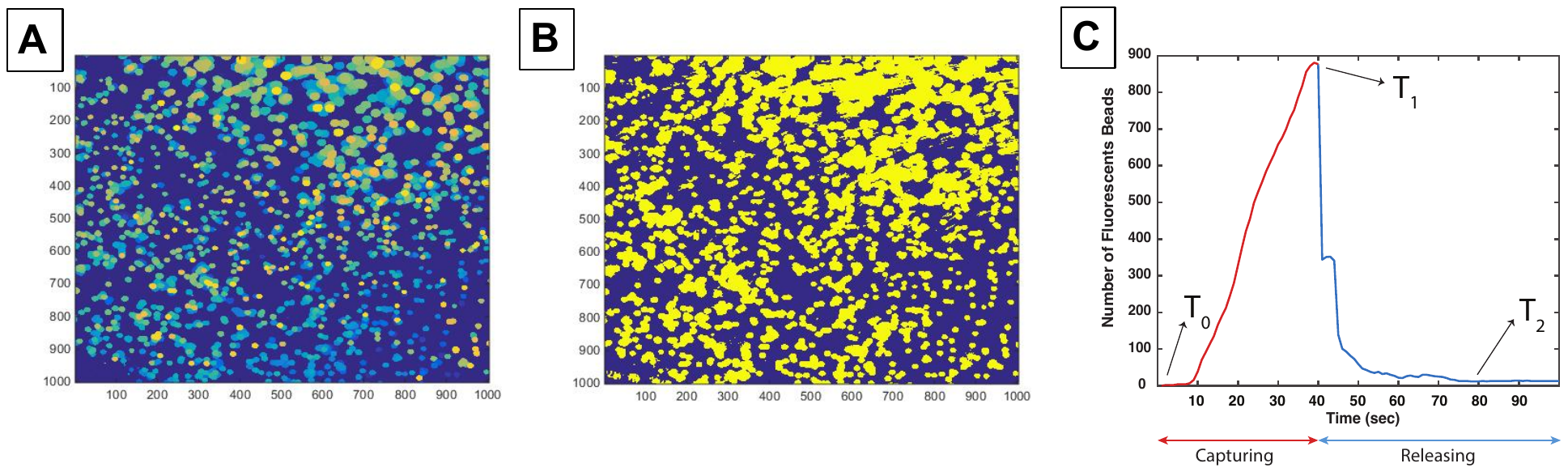

**Supplementary Figure 2; Counting of 10 micron fluorescent particles. A)** Residence time map **B)** Binarized residence time map **C)** Number of fluorescent particles - time plot showing capturing and releasing of micron particles over the experiment.

MATLAB code was generated to count the number of micron particles being captured and released over time. All the images in the time lapse series were binarized and summed over time to create the residence time map as it can be seen in figure S1-A. The color indicates the residence time of the particles on the membrane (where yellow indicates longer residence time). In order to ensure that floating particles were not counted as a captured particle, a threshold of 3 images was applied so that particles that were in the field of view for less than 3 images were excluded. Then, the images were binarized to yellow for the captured particles and blue colors for the membrane (figure S1-B). Number of particles was calculated by dividing the yellow colored area by the area of a single particle for every single image during the experiment (figure S1-C).

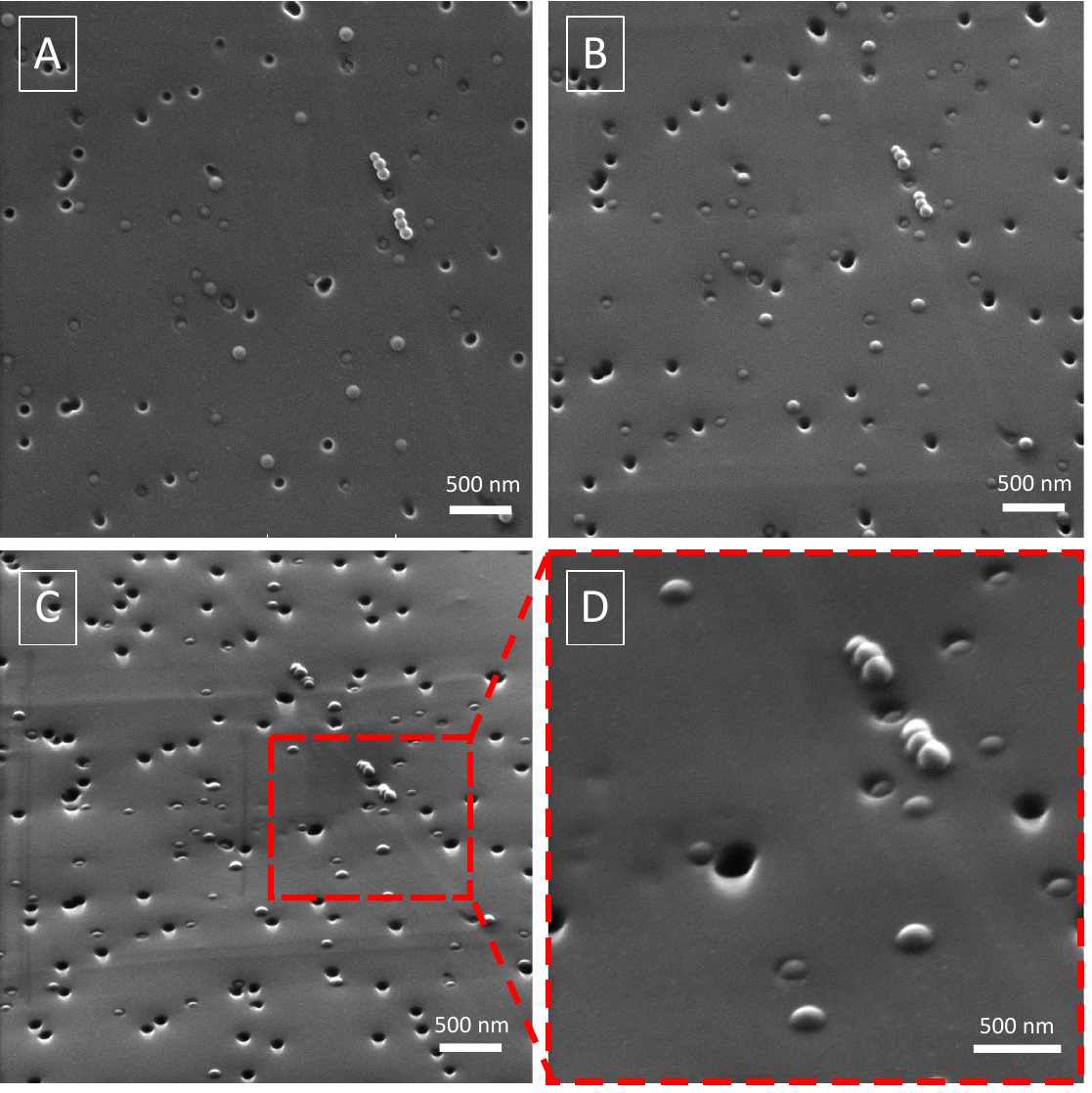

**Supplementary Figure 3; Characterization of nanoparticles capturing sites of TE membranes using SEM images.** Micrographs were taken at different stage positions to show different perspectives. **A)** Top down view **B)** 30° tilt **C)** 60° tilt **D)** High magnification image of 60° tilt sample.

SEM images after capturing-cleaning step suggested capturing of nanoparticles on the pores, inside the pore channels and on the surface of the membranes. In order to determine the capturing sites on the membranes, samples were gradually tilted and imaged as it can be seen in figure S3. High magnification SEM image of 60⁰ tilted sample showed captured particles on the pores, inside the pores, and on the surface of the membranes due to the charge interaction.

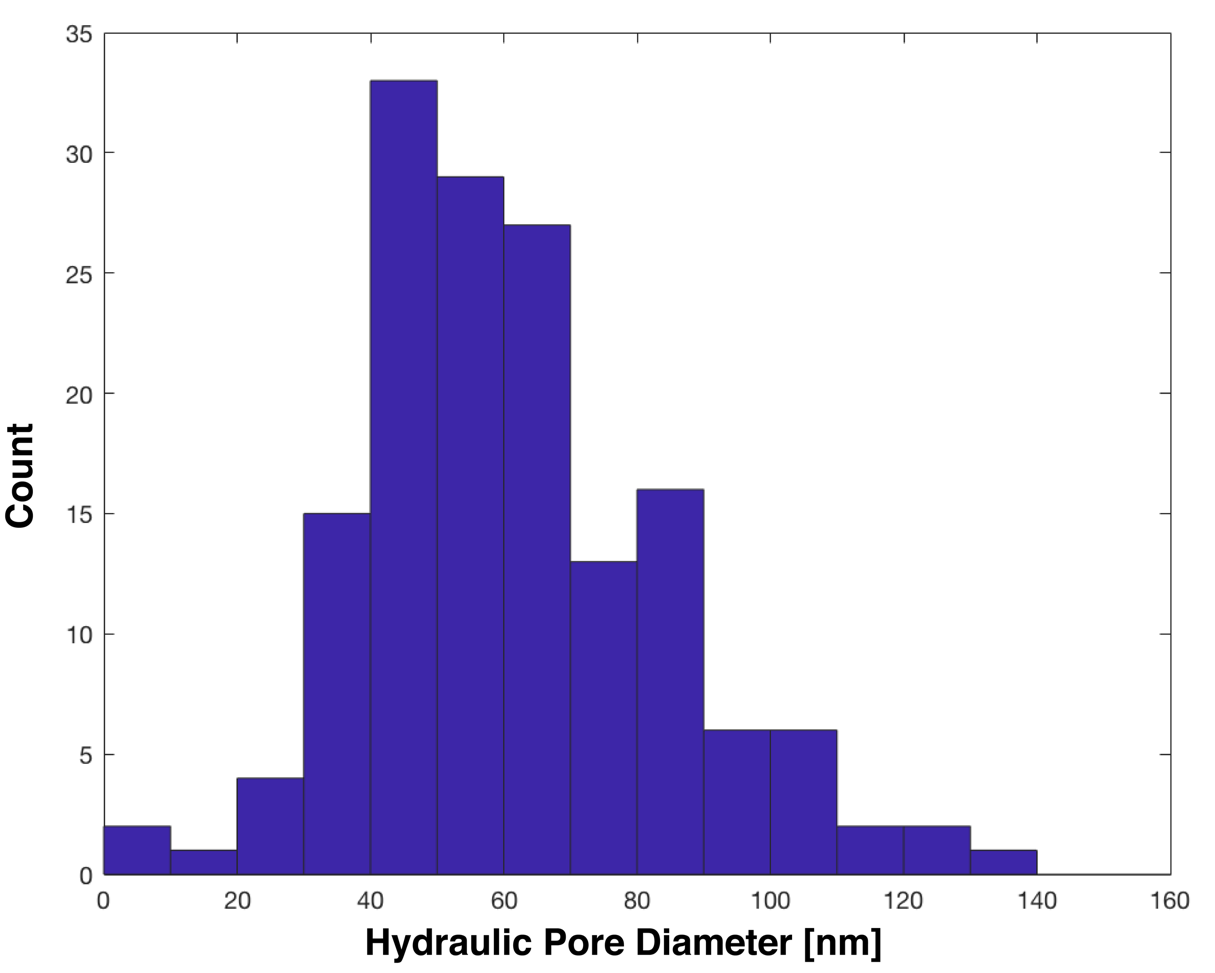

**Supplementary Figure 4: Pore Size Distribution for Nanoporous Silicon Nitride Membrane Chips**. The pore distribution of NPN chips is heterogeneous, but the median diameter is 80 nm. The etching process produces random, large pores, but these would improve the capture of a heterogeneous particle population such as EVs.

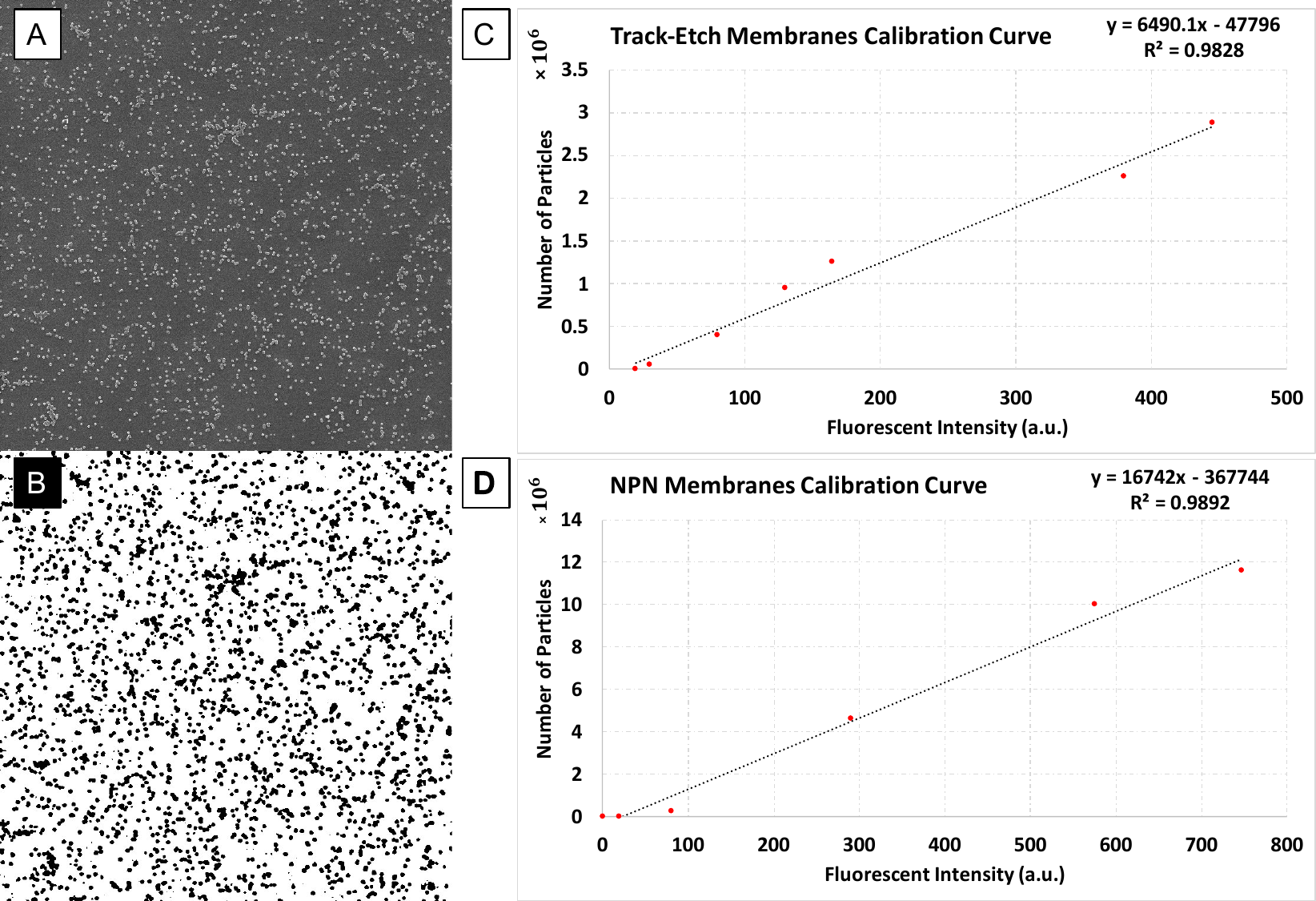

**Supplementary Figure 5; Calibration curves correlating the fluorescent intensity to the number of particles on TE and NPN membranes.** **A)** SEM image of a uniform deposition of fluorescent nanoparticles **B)** Binarized image **C)** TE membranes calibration curve **D)** NPN membranes calibration curve

Calibration curves for both PCTE and NPN membranes were obtained by correlating the fluorescent intensity with the number of fluorescent particles dried on the membrane. In order to avoid a coffee-ring effect during the drying process, water drops containing 100 nm monodispersed particles were dried on PCTE and NPN membranes at 80°C (Burkhart, Maki, & Schertzer, 2017). Drying the drop at higher degrees allowed the particles to accumulate at the air-liquid interface rather than at the drop edge leading to uniform deposition of particles (Y. Li, Yang, Li, & Song, 2016). The dried membranes were then placed back into the microfluidic devices and the channels were primed. Fluorescent images were taken using the Leica microscope and the intensity values were measured using ImageJ. Scanning electron microscopy was used to count the number of particles on the surface of the membranes. SEM images were binarized to black and white using ImageJ and number of particles in binarized SEM images was calculated by dividing the total area of particles by the area of a single particle. The best linear fit and the equation correlating the concentration of particles and the fluorescent signal were obtained and further used for analyzing the experimental data on PCTE and NPN.

**Supplementary Video 1; Layer by layer stacking process for fabrication of microfluidic devices.**

**Supplementary Video 2; Capturing and releasing of 10 micron fluorescent particles on 8 micron pore size PCTE membranes.**
